# Multiscale affinity maturation simulations to elicit broadly neutralizing antibodies against HIV

**DOI:** 10.1101/2021.09.01.458482

**Authors:** Kayla G. Sprenger, Simone Conti, Victor Ovchinnikov, Arup K. Chakraborty, Martin Karplus

**Author notes:** These authors contributed equally to this work (Kayla G. Sprenger, Simone Conti).

## Abstract

The design of vaccines against highly mutable pathogens, such as HIV and influenza, requires a detailed understanding of how the adaptive immune system responds to encountering multiple variant antigens (Ags). Here, we describe a multiscale model of B cell receptor (BCR) affinity maturation that employs actual BCR nucleotide sequences and treats BCR/Ag interactions in atomistic detail. We apply the model to simulate the maturation of a broadly neutralizing Ab (bnAb) against HIV. Starting from a germline precursor sequence of the VRC01 anti-HIV Ab, we simulate BCR evolution in response to different vaccination protocols and different Ags, which were previously designed by us. The simulation results provide qualitative guidelines for future vaccine design and reveal unique insights into bnAb evolution against the CD4 binding site of HIV. Our model makes possible direct comparisons of simulated BCR populations with results of deep sequencing data, which will be explored in future applications.

Author Summary

Vaccination has saved more lives than any other medical procedure, and the impending end of the COVID-19 pandemic is also due to the rapid development of highly efficacious vaccines. But, we do not have robust ways to develop vaccines against highly mutable pathogens. For example, there is no effective vaccine against HIV, and a universal vaccine against diverse strains of influenza is also not available. The development of immunization strategies to elicit antibodies that can neutralize diverse strains of highly mutable pathogens (so-called ‘broadly neutralizing antibodies’, or bnAbs) would enable the design of universal vaccines against such pathogens, as well as other viruses that may emerge in the future. In this paper, we present an agent-based model of affinity maturation – the Darwinian process by which antibodies evolve against a pathogen – that, for the first time, enables the *in silico* investigation of real germline nucleotide sequences of antibodies known to evolve into potent bnAbs, evolving against real amino acid sequences of HIV-based vaccine-candidate proteins. Our results provide new insights into bnAb evolution against HIV, and can be used to qualitatively guide the future design of vaccines against highly mutable pathogens.

## Introduction

Although great progress has been made in the development of therapeutics and annual vaccines against diseases such as HIV and influenza, respectively, the need for universal vaccines for these diseases remains high. Approximately 38 million people are still living with AIDS today [1], and up to 650,000 annual deaths globally are still attributed to influenza [2]. Unfortunately, such viruses undergo rapid mutations that can render the host immune response ineffective, greatly reducing vaccine efficacy. Special classes of antibodies (Abs) called “broadly neutralizing antibodies”, or bnAbs, have now been discovered that are effective against HIV [3] and influenza [4]. However, it remains unclear how to elicit them via vaccination. This is largely due to an incomplete understanding of how the immune system responds to encountering multiple variant Ags, a situation that can arise during infection with a rapidly evolving pathogen.

B cell receptors (BCRs, or equivalently Abs) evolve through the evolutionary process of affinity maturation (AM) [5], which occurs within Germinal Centers (GCs). The large diversity of the human germline BCR repertoire implies that, after encountering an Ag, some BCRs will bind to the Ag, albeit weakly. If the binding affinity is above a certain threshold, those B cells will get activated, undergo replication inside the GC, and accumulate mutations in their BCRs. The resultant mutated B cells are continuously recycled and selected on the basis of their affinity for the Ag [5].

In the presence of multiple Ags, which can be administered simultaneously or sequentially in a vaccine or be encountered naturally as mutated versions of the original infecting virus, B cells evolve to recognize each of the Ags to some extent. If the Ag binding sites (epitopes) presented to the immune system are sufficiently dissimilar, the activated memory B cells will experience conflicting selection forces, which have been described by a “frustration” parameter in our past work [6–8]. Changing the degree of dissimilarity between the Ags administered in a vaccine has been used to modulate the level of frustration imposed on GC reactions, and can have the effect of focusing BCR responses on conserved Ag residues, leading to the elicitation of bnAbs [6–8].

Past computational studies have identified additional variables that influence the evolution of BCRs, including Ag concentration [6, 8] and the temporal pattern of Ag administration [7, 8]. However, it remains unclear how to set these variables to optimize bnAb production. Additionally, due to the coarse-graining used in past computational models of AM [6–15], most of these models cannot make predictions about the actual BCR and Ag sequences that will best promote bnAb formation during AM.

Several computational studies employed approximate AM schemes, in which Ab structures are used in protein docking, undergo multiple rounds of *in silico* mutagenesis, followed by evaluation of binding free energy changes using force field-based scoring functions [16–18]. These studies use nucleotide sequences to represent the BCRs, but do not explicitly include important aspects of AM such as clonal competition and interference. Additionally, mutations are introduced into the BCRs on an *ad hoc* basis, neglecting the mutational biases of activation-induced cytidine deaminase (AID [19]), the enzyme that induces mutations during AM. Therefore, such approaches are unlikely to serve as reliable guides for studying the effects of vaccines on BCR/Ab evolution.

In this study, we describe a computational model of AM that (1) utilizes actual nucleotide sequences of BCRs and Ags, (2) employs an experimentally-informed model [19] for the identity and location of BCR nucleotide mutations introduced during AM, and (3) incorporates an efficient method to calculate BCR/Ag binding free energies. We believe this to be the most realistic model of AM currently available, which has the capacity to guide vaccine design efforts against highly mutable pathogens such as HIV and influenza.

As an application of the model, we study the evolution of the germline precursor of VRC01, a potent bnAb against the CD4 binding site (CD4bs) of HIV, in response to vaccination with multiple rationally-designed [20] HIV-based Ags administered in different temporal patterns. As in our past work [8], we assume that the desired germline B cells have already been activated [21]. Our results show that administering Ags in a temporal pattern that continuously increases the amount of imposed frustration maximizes the mean breadth of the resultant BCRs. We also find that administering certain Ags leads BCRs to evolve mutations that increase their degree of interfacial amino acid composition/electrostatic pattern matching against the CD4bs. We propose that this is an important mechanism by which maturing BCRs develop high affinity for the CD4bs of HIV.

## Results

### Simulated affinity maturation

A number of coarse-grained models to simulate AM are in the literature, many of which were developed within the past five years [6–15]. These models use coarse-grained representations for the BCR and Ag sequences, without a clear connection to the constituent nucleotides or corresponding amino acids. Here, we started from a coarse-grained model [8] and introduced actual nucleotide and amino acid representations for the BCRs and Ags, respectively. The model starts by seeding the GC with known germline BCR sequences (see Methods for details). We focus on HIV, and therefore seed the GC with the VRC01 germline [21] (VRC01GL) sequence, which is known to evolve into the potent bnAb VRC01. The founding B cell replicates without mutation or selection until the population reaches a size of 128 cells [22]. Then, the B cells start to accumulate mutations in their BCRs according to an AID-based mutation model [19], after which they compete for Ag and T cell help; 70% of the surviving B cells are selected randomly to undergo a new cycle of mutation and selection, and 30% exit as memory B cells [23]. The simulation ends when the B cell population recovers its initial size (successful outcome), or when all B cells die (unsuccessful outcome). In subsequent immunizations, the GC is seeded with three sequences randomly sampled from the memory BCR population produced during the previous immunization. An overview of the model is shown in Fig. 1.

**Fig. 1.**
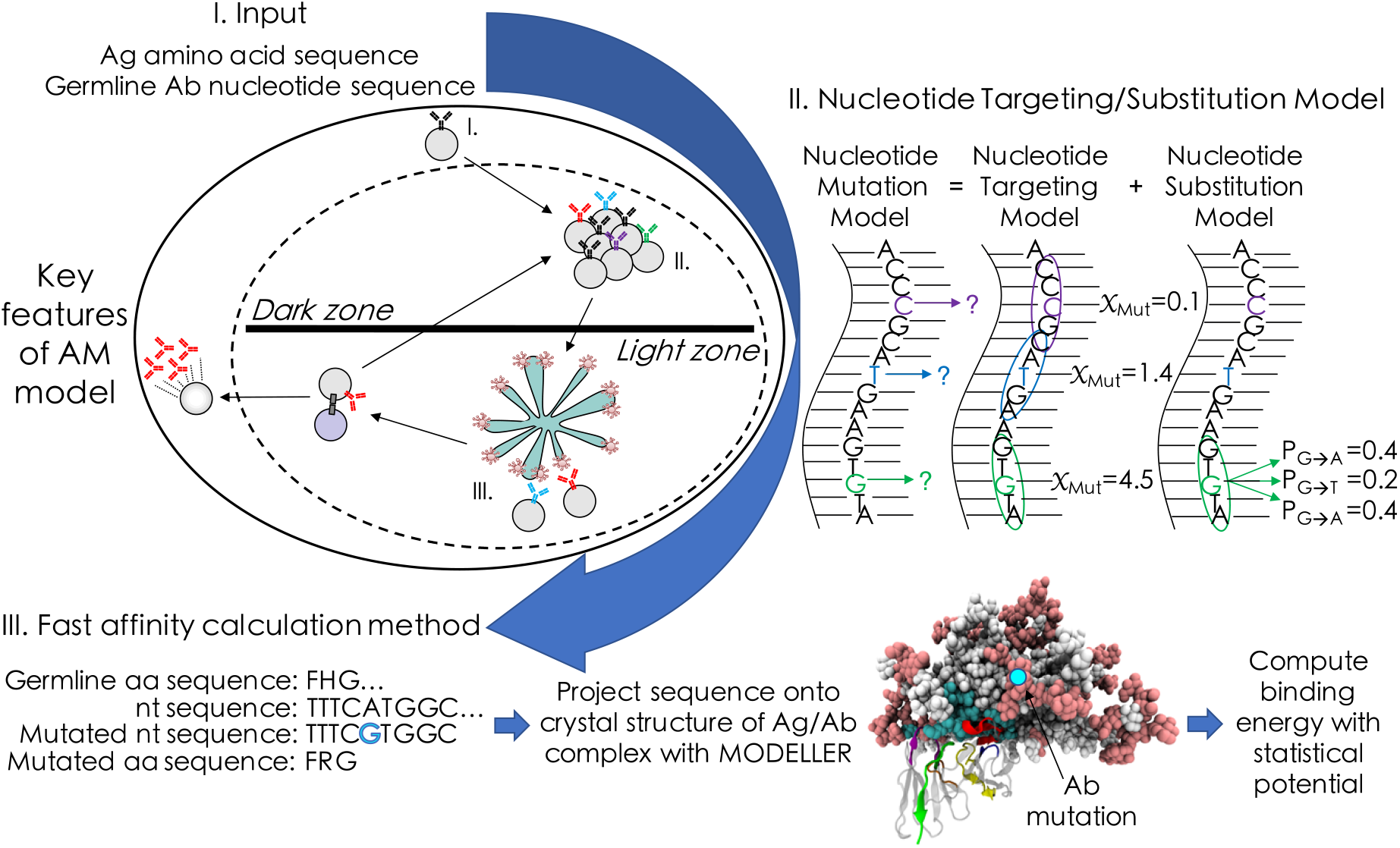
Simulation framework for the AM model, highlighting new features. Amino acid (aa) and nucleotide (nt) sequences of the Ag and germline BCR, respectively, are input into the model in step I. During B cell proliferation and mutation (step II), a mutation model [19] is used to determine where and which mutations will occur in the BCR sequences; nts that have a higher mutability score (*χ*_Mut_) – which accounts for the effects of surrounding nts – have a greater chance of being selected for mutation. This is followed in step III by BCR/Ag binding free energy calculations, which utilize the program Modeller [25] and published statistical potentials [26]. Final steps in the model include a second selection step, recycling, and differentiation (details in text). BCR breadth is computed as the fraction of Ags in a panel for which the BCR binds above a chosen threshold (see Methods).

To reduce the computational complexity, we limited mutations to the complementarity-determining regions (CDRs) of the BCRs, and we did not introduce framework mutations, which are more likely to affect BCR stability [24]. Lastly, to compute the BCR/Ag binding free energy, we first create atomic models of the BCR/Ag complex with Modeller [25], and then evaluate the binding free energy using scoring functions from the literature [26]. On a standard 16-core CPU, each binding free energy calculation takes approximately one minute, rendering the AM simulations – which can require upwards of 15,000 calculations/GC – feasible. Overall, for the production runs and analysis carried out in this work, we computed nearly one million binding free energies for unique BCR/Ag complexes.

### Temporal pattern of Ag administration influences the breadth of the BCRs

We studied the outcomes of AM in response to 36 sequential vaccination protocols (Table S1 in the SI). The protocols are composed of a maximum of three sequential immunizations of four possible Ags, namely the wildtype BG505 SOSIP (WT) and three BG505-based derivatives designed to maximally promote the bnAb evolution against the CD4 binding site (CD4bs) of HIV [20]. These three designed Ags are identified as KR, EU, and HQ (see Methods). The concentration profile was kept the same across all vaccination protocols (see Methods); therefore, differences in simulation outcomes are due solely to differences in Ag sequences or temporal patterns of administration.

Upon a single immunization with each of the four Ags, we find that the mean BCR breadth increases as follows: WT (breadth = 0.00) < KR (breadth = 0.005) < EU (breadth = 0.11) < HQ (breadth = 0.15). These results show that the choice of administered Ag has a significant influence on breadth. We next investigated whether the mean BCR breadth could be optimized by administering these Ags in different temporal patterns (Fig. 2A). Averaging the results across all 1-, 2-, or 3-Ag protocols, we find that the mean BCR breadth increases with the number of immunizations (breadth = 0.07, 0.39, and 0.73 for 1-, 2-, and 3-Ag protocols, respectively). However, the mean breadth is also observed to vary widely across the 2- or 3-Ag protocols, which shows that the temporal Ag administration pattern can have a significant impact on BCR breadth. We also find that the mean number of GC cycles is correlated with breadth (Fig. 2B), as more GC cycles afford BCRs more time to accumulate mutations that confer breadth.

**Fig. 2.**
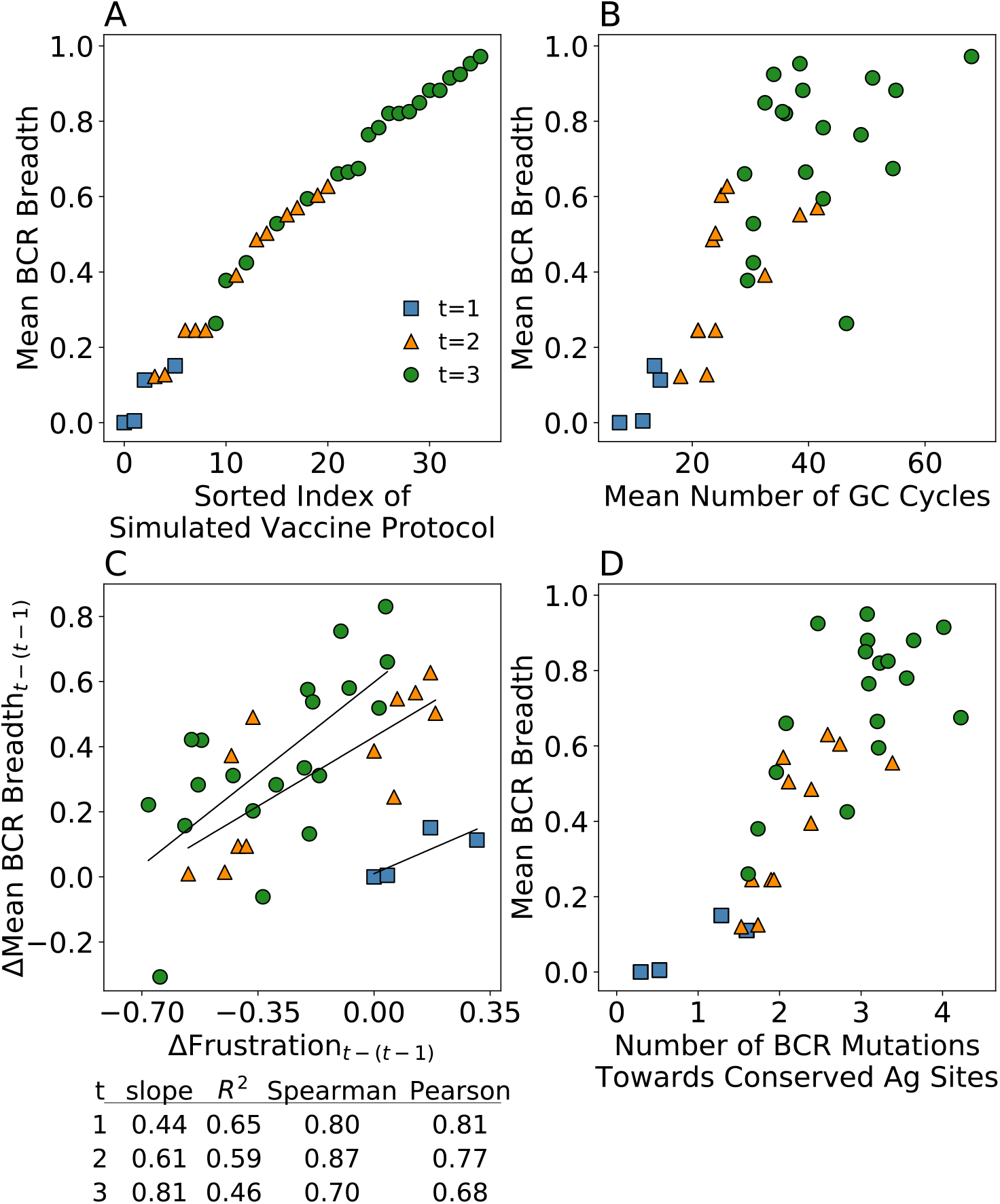
Results from AM simulations. (A) mean BCR breadth of vaccination protocols with t=1, 2, and 3 single-Ag sequential immunizations, (B) mean BCR breadth vs. mean number of GC cycles, (C) changes in mean BCR breadth vs. changes in frustration between current (t) and former (t-1) immunizations, and (D) mean BCR breadth vs. mean number of mutations towards conserved Ag residues. Solid lines in (C) represent the linear fits used to calculate the R^2^ values presented in the table below the graph. Error bars represent the standard deviation of two independent simulations of n=1,000 GC reactions (error bars for mean BCR breadth are only shown in (A) for clarity).

To understand the effects on AM of changing the temporal Ag administration pattern, we calculated the frustration imposed on the GC reactions by the different vaccination protocols, which was defined as the opposite sign of the average binding free energy of the GC-seeding BCR(s) and the Ag administered in the subsequent immunization. As such, the “frustration” is really a metric of the extent to which the existing B cell population is thrown off of a previous steady state. The change in imposed frustration versus the change in mean BCR breadth between the current and former immunization is shown in Fig. 2C. We find a positive correlation between the change in frustration and in the mean BCR breadth, with the increase being higher in each subsequent immunization (slopes reported in Fig. 2C). In addition, with the current set of four Ags, we find that progressively fewer protocols result in actual increases in frustration (i.e., rather than smaller decreases; those to the right of zero on the x-axis) with each additional immunization. This could indicate that three sequential immunizations with these Ags are not necessary to elicit sufficiently high-breadth Abs. Mechanistically, this is because BCRs become increasingly capable of compensating for increasing frustration using strengthened interactions with conserved Ag residues. We demonstrated this by calculating the mean number of BCR mutations towards Ag residues that are considered to be conserved in the CD4bs [20] (see Methods and Table S3 in the SI). Fig. 2D shows that the mean number of BCR mutations towards conserved Ag residues increases from 0.95 to 2.2 to 3.0 after one, two, and three immunizations, respectively. Consequently, we observe a strong positive correlation (R^2^=0.79) between the number of conserved site mutations and the mean BCR breadth.

In summary, we find that the elicitation of bnAbs is strongly dependent on the temporal pattern of Ag administration. Specifically, we find that vaccination protocols that impose the greatest increases in frustration from one immunization to the next result in the greatest increases in breadth. The increased frustration results in longer GC reaction times, which allow BCRs more time to acquire mutations that confer breadth. These results are consistent with our past studies with coarse-grained models. [8, 27]

### BnAbs employ interfacial composition and electrostatic pattern matching (ICM) to bind to the CD4bs of HIV

To provide context for our simulation results, we performed a detailed analysis of the interfacial mutations that VRC01GL evolved *in vivo* during its maturation into its bnAb form, VRC01. Analyzing crystal structures of VRC01GL [21] and VRC01 [28] in complex with HIV-based Ags, 26 and 29 interfacial Ab residues, respectively, were identified to be within 4Å of the Ag. For both sets of residues, we determined the fraction of polar, apolar, acidic, and basic amino acids (Fig. 3A, right; see Methods). For the antigen, the same coarse-graining procedure was carried out on the interfacial residues of 106 HIV Ag sequences from the Seaman panel [29]. The results were then averaged across the panel (Fig. 3A, left; see Methods), in order to account for the highly mutable nature of HIV.

**Fig. 3.**
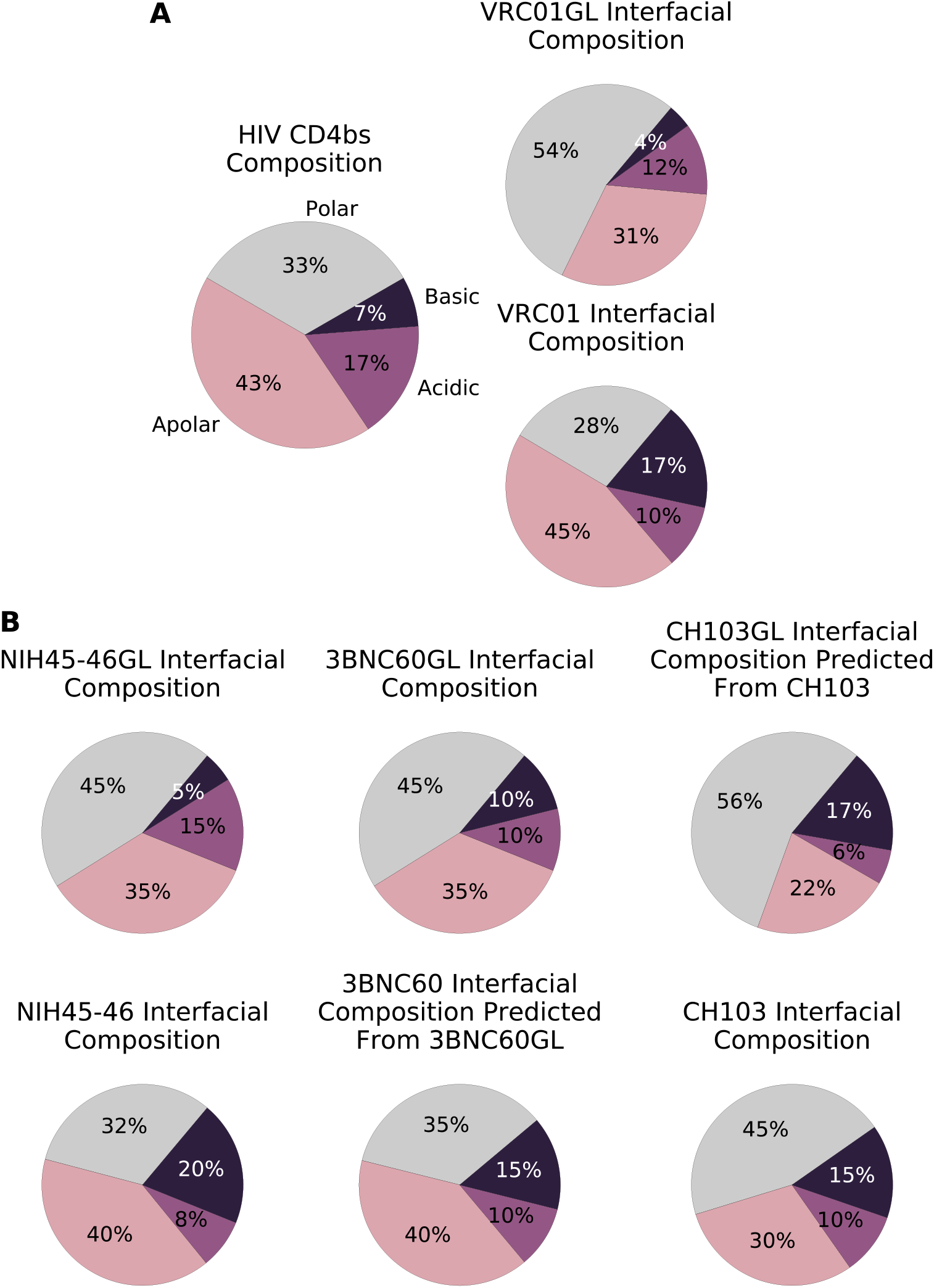
Anti-CD4bs bnAbs employ interfacial composition and electrostatic pattern matching (ICM) during AM. (A) Interfacial amino acid compositions of the CD4bs of HIV (left), VRC01GL (top right), and VRC01 (bottom right). (B) Interfacial amino acid compositions of germline Abs NIH45-46GL (top left), 3BNC60GL (top middle), and CH103GL (top right), and their bnAb counterparts (bottom row).

Before AM, VRC01GL had significantly higher and lower fractions of polar and apolar interfacial residues than the CD4bs (Fig. 3A). However, in the case of the affinity-matured Ab, VRC01, these two categories match much more closely those of the CD4bs. Moreover, the basic interfacial fraction increases from 4% in VRC01GL to 17% in VRC01, to match the acidic fraction of the CD4bs at 17%, and the acidic interfacial fraction decreases from 12% in VRC01GL to 10% in VRC01, to better match the basic fraction of the CD4bs at 7%. Overall, a trend towards interfacial composition and electrostatic pattern matching (ICM) was observed between the evolving VRC01 Ab and the Ag binding site.

To validate these results, this analysis was performed on three other bnAbs targeting the CD4bs, for which both the germline and mature sequences were available: NIH45-46 [30, 31], 3BNC60 [30], and CH103 [32] (see Methods). Fig. 3B shows that these additional Abs also employed ICM during their maturation. This is particularly interesting for CH103, which binds to the CD4bs in a significantly different pose than the other bnAbs we considered (Fig. S1). Overall, these results imply that ICM is likely to be a general mechanism employed during AM and is an observable that can be monitored in the simulations. In the SI, we explore the biological driving forces behind these results, which include differences in the mutability of the codons encoding different amino acid types at the interface, and the number of nucleotide mutations required to transition from one amino acid type to another.

The degree of ICM of the *in silico-*produced BCRs was quantified using a chi-squared statistic χ^2^-based formula on 22 interfacial residues in each BCR sequence (see Methods). The ICM score measures the summed deviation of the interfacial fraction of each amino acid type from its value in the CD4bs of HIV (n.b., the fraction of acidic residues is compared to the fraction of basic residues in the CD4bs, and vice versa; see Eqn. 3, 4 in Methods), subtracted from unity. The ICM scores calculated for VRC01GL and VRC01 are 0.73 and 0.99, respectively. Computing the mean ICM score for the 36 vaccination protocols, we find that among the protocols with the same first administered Ag, the resultant BCRs tend to increase their ICM scores with each subsequent immunization (Fig. 4, Fig. S4), in line with the biological trends observed for the four bnAbs discussed above. Among protocols starting with each of the four Ags, those protocols in which the first administered Ag is EU, WT, or HQ consistently resulted in relatively low, intermediate, and high degrees of ICM, respectively. Protocols in which the first Ag was KR resulted in a wide range of ICM scores, implying that KR is overall a weak driver of ICM, despite the high average ICM score of the KR-KR protocol (0.92). Interestingly, we observe only a weak correlation between the degree of ICM and the mean breadth of the BCRs produced in the simulations (R^2^=0.17), though this correlation improves slightly upon reducing the stringency of the BCR breadth threshold (Fig. S3).

**Fig. 4.**
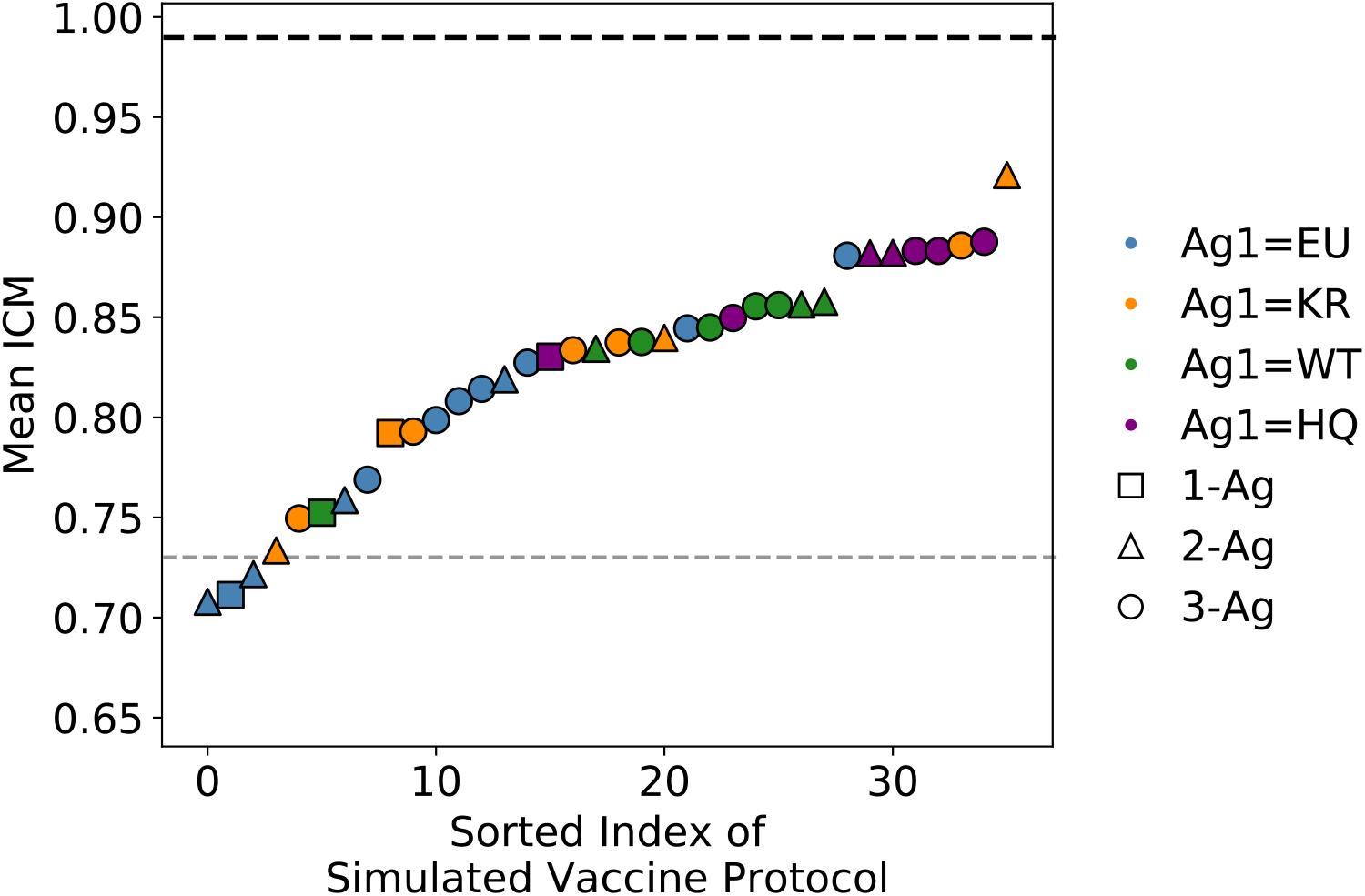
Results from AM simulations: mean weighted degree of interfacial composition and electrostatic pattern matching (ICM) of all 1-, 2-, or 3-Ag protocols, sorted by the mean ICM score. Error bars represent the standard deviation of two independent simulations of n=1,000 GC reactions. Gray and black dotted lines indicate the respective values for VRC01GL and VRC01.

To understand why some Ags elicit BCR populations with higher ICM scores over others, we decomposed the ICM scores in Fig. 4 into their individual components (i.e., changes in the interfacial fraction of each amino acid type in the BCR population over time). These results and the corresponding analysis are provided in the SI (Fig. S4). The results show clear trends between the types of BCR mutations made and the corresponding ICM scores. For example, protocols beginning with the EU Ag typically resulted in acidic-to-polar mutations, leading to poor ICM scores. Conversely, protocols beginning with the HQ Ag typically resulted in polar-to-basic mutations, resulting in high ICM scores.

## Discussion

Conventional vaccination approaches to evolve antibodies (Abs) against pathogens such as HIV and influenza are largely unsuccessful due to high mutability or antigenic drift. Broadly neutralizing Abs (bnAbs) could overcome this challenge, though it remains unclear how to elicit them via vaccination. Consequently, research focus is shifting towards identifying vaccine design variables that can facilitate bnAb evolution during affinity maturation (AM); e.g., sequences and concentrations of the administered antigens (Ags) [6,7,33–39]. Previous computational studies utilized coarse-grained representations for the B cell receptors (BCRs) and Ags. In this work, we developed a multiscale computational framework that models the AM of actual BCR sequences in response to vaccination with atomic-resolution Ags. This model enables future comparisons between simulations and experimental results from, e.g., deep sequencing of BCR repertoires.

Changing the Ags and their concentrations during vaccination disrupts the normal progression of AM by perturbing the memory BCR population, imposing what we termed “frustration” as in our past work [6–8]. Here, we manipulated the level of frustration by administering four variant Ags, three of which were explicitly designed to improve the breadth of the elicited anti-HIV Abs [20], in 36 different temporal patterns. Our results show that increasing the imposed frustration increases the number of GC cycles during which BCRs can accumulate breadth-conferring mutations. Increasing frustration counteracts the fact that BCR interactions with conserved Ag sites are strengthened with each subsequent immunization, effectively lowering the frustration. This finding is qualitatively consistent with results from our previous work with coarse-grained models [8, 27] that showed increasing the frustration upon the second immunization imposed a stronger selection force on activated memory B cells to evolve additional mutations towards conserved Ag residues.

The above comparison is only qualitative because many more simulations could be performed with the coarse-grained model due to the much lower computational cost, enabling conclusions about B cell survival rates. However, if trends from our past work hold true here as well, they would imply that rather than attempting to maximize the level of frustration in each immunization, there is an optimally increased level of frustration to impose (i.e., Ag sequence to use, in this case) in each immunization. Such an approach is predicted to simultaneously optimize the production of high-breadth Abs and B cell survival rates [8].

An important finding from this work, which was not possible to obtain with previous coarse-grained models, is that the VRC01 bnAb evolved mutations at the interface that enabled interfacial composition/electrostatic pattern matching against the CD4bs itself. We showed that three other anti-CD4bs bnAbs also employed this mechanism – which we term ICM – during their evolution. Results from the AM simulations show that the produced BCRs also increase their degree of ICM over time, though the final degree of ICM achieved by the BCRs appears to depend strongly on the first Ag that they encounter. After immunizing with the first Ag, B cells are selected for mutations that improve their binding affinity to that particular Ag. The effect of these mutations on the degree of ICM varies widely depending on the Ag, in part because of the many possible affinity-increasing mutations. This effect is likely diminished for anti-HIV bnAbs due to the numerous strains of the virus they encounter over their many years of maturation.

We discuss various driving forces behind ICM in the SI. In addition, better ICM may enable BCRs to more closely approach an Ag, thus allowing contacts with the conserved residues of the Ag. This, in turn, may increase the selection force to evolve mutations that bind better to the conserved residues, thus conferring breadth. Such a mechanism may promote bnAb evolution even though ICM per se does not correlate highly with breadth. Also, it remains unknown whether potent (but not necessarily broad) antibodies increase their degree of ICM during maturation. This would indicate that ICM is a more general feature of antibody evolution, a pattern matching mechanism which allows for general compatibility in the interactions between BCRs and Ags, before precise mutations are made to enable BCR binding to conserved Ag residues. Future longitudinal studies of bnAb evolution may help to shed light on this factor.

In view of our results, we suggest that a promising route to a universal vaccine for HIV is to first administer an Ag that promotes BCR mutations which increase the degree of ICM against the CD4bs (although we acknowledge that designing such immunogens may be nontrivial). This step should be followed by sequential immunizations with Ags that have an optimal number of increasingly different amino acids in the variable residues surrounding the conserved residues to achieve an optimally increasing temporal frustration profile. This procedure is expected to optimize the breadth of the mature BCR/Ab population, and according to our past work [8], it will also maximize the produced bnAb titers.

It is of high importance to understand how differences in Ag sequences lead to differences in the types of evolved BCR mutations. An advantage of our model is the ability to gain such molecular level insight by studying in detail the BCR/Ag complexes produced at various stages of AM. This would enable true Ag sequence design, future experimental testing of which would provide the ultimate assessment of how realistic our free energy estimator is for guiding the choice of actual Ags. While beyond the scope of the current work, such analyses and design studies are planned for the future.

## Methods

We first describe the multiscale model used to simulate the affinity maturation (AM) of a B cell population in a germinal center (GC) in response to vaccination. Following this, details of the analysis protocols are presented.

### Seeding of the germinal center

We assume that a germline (GL)-targeting scheme has occurred prior to vaccination to activate the desired bnAb precursor B cells, in the present case those of the VRC01 class [21]. Experimentally, a few to a hundred B cells have been shown to seed a GC [40]. Due to the sparsity of data currently available of VRC01 class precursor B cells bound to HIV-based antigens (Ags), we modeled the GCs initiated upon the first vaccine immunization as being seeded with a single VRC01 GL precursor B cell [21]. The nucleotide sequence of the B cell receptor (BCR) is input directly into the model, along with the amino acid sequence of the antigen(s) (Ag(s)) used for vaccination.

### Antigen sequences

Four different Ag sequences were employed in the *in silico* vaccination schemes. These include three Ags (identified as KR, HQ and EU), and the sequence of the wildtype HIV BG505 SOSIP, which is referred to as WT. The three Ags KR, HQ and EU have been previously designed to have a high potential of eliciting bnAbs to the CD4bs when used in an HIV vaccine [20]. These are based on the WT sequence, with selected mutations introduced from the naturally-occurring Ags KR423280, HQ217523, and EU577271; see original reference for details on the design [20].

### B cell expansion and mutations in the dark zone

After the GC has been seeded, the B cell population undergoes pure expansion (i.e., no mutation/selection) in what is known as the “dark zone” for approximately one week, to reach a population size of 128 cells in the simulation (2^7^). The gene encoding the enzyme activation-induced cytidine deaminase (AID) is then activated, facilitating the accumulation of mutations in the BCRs at a rate of roughly 0.14 mutations per sequence per division [41], with each B cell dividing on average twice per GC cycle [42]. As in our previous work [24], we set the probability of a mutation occurring in the complementarity-determining regions (CDRs) to 0.85 and in the framework regions (FRWs) to 0.15. However, to reduce the cost of the computational modeling, only CDR mutations were considered here to have an affect the BCR/Ag binding free energy. This was also necessary because our scoring function currently does not account for the additional complexities of many FRW mutations (e.g., effects on protein stability [24]). We also assume that 30% of all mutations are lethal to more closely mimic what has been shown to be the case in experiments [41].

### Mutation model

Mutations in the BCR sequences are introduced by AID on the nucleotide level. Where and which mutations arise in the BCRs during each cycle of the GC reaction is largely a function of the dynamics of AID, which shows pronounced biases towards certain patterns or groups of nucleotides [19]. The tendency of a particular nucleotide in a BCR sequence to be mutated during AM depends primarily on the surrounding microsequence environment of that nucleotide; specifically, it has been shown to depend on the identities of the two neighboring nucleotides on either side in the sequence. [19] Yaari *et al.* [19] considered all possible combinations of such groups of five nucleotides, which they deemed “5-mers”, and determined the relative tendency (“mutability score”) of each 5-mer to be targeted for mutation by comparing the sequences of over 1 million Abs before and after AM [19]. In our model, for a given BCR sequence in a given GC cycle, we first determine all of the relevant 5-mers in a sliding window and look up their corresponding mutability scores. Next, we normalize this subset of mutability scores to convert them into probabilities. They are then used to determine the single nucleotide that will be mutated in the BCR sequence during that GC cycle. In a similar manner as described above, Yaari *et al.* also determined the probability that the central nucleotide in each 5-mer is mutated to each of the other nucleotides (i.e., to A, C, G, or T) [19]. After determining the single nucleotide to undergo mutation in a given BCR/GC cycle, these substitution probabilities are then used to sample the nucleotide.

### Selection, recycling, and differentiation

After acquiring mutations in their receptors, B cells compete with other B cells for binding to the Ags and receiving support from T helper cells [22]. B cells whose receptors are able to bind more strongly to the Ags are in turn able to consume, break down, and display more Ags on their surface, which ultimately increases their chance of receiving productive proliferation signals from T helper cells [22]. B cells that internalize low levels of Ags do not interact productively with T helper cells and undergo apoptosis [43–45]. As in our past work [8], this behavior is modeled by Equation 1:

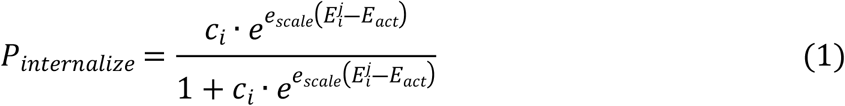

where *c*_*i*_ is the concentration of Ag *i*, 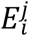 is the binding free energy between Ag *i* and B cell *j*, *E*_*act*_ is the activation energy, and *e*_*scale*_ is a pseudo inverse temperature. The activation energy for seeding a GC was chosen to be equal to the weakest binding free energy between the four Ags and VRC01GL – that of EU at -9.01 kcal/mol – such that all four Ags could induce an effective immune response. The value of *e*_*scale*_ (5.8 (kcal/mol)^-1^), along with the value of the breadth threshold (-9.9 kcal/mol; see below), were adjusted such that a single immunization with any of the Ags would produce mostly low-breadth Abs (a breadth ≤ ∼0.15). The above condition mimics the fact that breadth is unlikely to result from a single immunization. This is demonstrated by the finding that in natural infection, bnAbs typically require several years to evolve, and that in vaccination studies with mice, they emerge only after several sequential immunizations with variant Ags [7, 33]. How the Ag concentrations, *c*_*i*_, used in the simulated immunization schemes were chosen and how the BCR/Ag binding free energies, 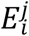, were calculated are described in the following sections.

B cells that survive this first selection test continue on to compete for T cell help; the top 70% of binders are selected for T cell help. From this pool of B cells that are positively selected, 70% are then chosen randomly to be recycled back to the dark zone for additional rounds of mutation and selection to iteratively increase their affinity for the Ag [23]. The other 30% of B cells exit the GC to simulate differentiation into plasma and memory cells [23].

### Antigen concentration profiles

The Ag concentration used in each successive immunization (c_1_=0.25, c_2_=0.05, c_3_=0.01) was chosen through trial-and-error to maximize the number of protocols for which we could obtain successful results (i.e., GC reactions that terminated due to recovery of the B cell population to its initial size of 128 B cells). In total, 48 protocols were initially considered: 4x3x2 unique 3-Ag protocols, 4x3 unique 2-Ag protocols, and 12 same-Ag protocols (4 3-Ag, 4 2-Ag protocols, and 4 single-Ag protocols). At the final chosen concentrations, we were able to obtain results for 36 of the 48 protocols; for the other 12 protocols, GC reactions did not terminate successfully. For example, after first immunizing with either WT or KR, B cell populations decreased rapidly, and did not recover after a second immunization with EU. Thus, we could not obtain results for the following 6 protocols: WT-EU, WT-EU-KR, WT-EU-HQ, KR-EU, KR-EU-WT, or KR-EU-HQ. Also, after first immunizing with HQ, the resultant B cell populations had matured sufficiently such that the B cell populations did not undergo any AM after subsequent administration of WT or HQ. Thus, we also could not obtain results for the following 5 protocols: HQ-WT, HQ-WT-KR, HQ-WT-EU, HQ-HQ, or HQ-HQ-HQ, and for WT-KR-HQ for similar reasons.

### Binding free energy and frustration calculations

BCR/Ag binding free energies are computed on-the-fly during the AM simulations, enabled by an efficient pipeline of steps that make use of several pre-existing external resources. First, we convert the mutated nucleotide BCR sequences into amino acid sequences. We then use Modeller [25] to graft these mutated sequences and that of the particular Ag used for immunization onto the crystal structure of VRC01 in complex with an HIV-based Ag [28]. Twelve complexes are independently built with Modeller for each calculation to obtain reliable average estimates for the BCR/Ag binding free energy. Following this conversion of sequence to structure, the Rykunov-Fiser statistical pair potential [26] is used to compute the binding free energy; see the SI for more details. The resulting protocol shows a Pearson correlation coefficient with experimental binding affinity data of 0.74 and an RMSE of 0.59 kcal/mol. Although approximate, this protocol allows for the computation of one binding free energy in about one minute using a 16-core processor. For the set of simulations performed in this study, the above protocol was used to calculate the binding free energy of nearly 1 million unique BCR/Ag complexes.

The calculated binding free energies were used directly in calculations of the level of frustration imposed on the GC reactions due to vaccination; since we did not change the concentration profiles across the protocols, changes in frustration could be linked directly to changes in Ag sequence or temporal pattern of administration. The frustration of a given immunization was defined as the opposite sign of the average binding free energy of the GC-seeding BCR(s) (germline BCR for 1-Ag protocols; memory BCRs for multi-Ag protocols) against the next-administered Ag, since stronger BCR/Ag binding free energies imposes less frustration by increasing the probability of internalizing Ag (see Eqn. 1 in the Methods). The change in frustration, Δ*F* was then computed between the current and former immunizations (Δ*F*_*t*=3―*t*=2_, Δ*F*_*t*=2―*t*=1_, and Δ*F*_*t*=1―*t*=0_). With regard to the 1-Ag protocols, ‘former’ refers to a reference immunization with frustration set equal to the weakest binding free energy between VRC01GL and the four Ags (that of VRC01GL and WT, at -9.32 kcal/mol), multiplied by -1.

### GC reaction termination

The GC reaction terminates if either: (a) the B cell population recovers to its initial size of 128 B cells in the first immunization or 384 B cells in subsequent immunizations (see next section) – termed a successful reaction – or (b) if all of the B cells die. Simulations were rerun until two successful trials of each protocol were achieved.

### GC reseeding with memory B cells

When multiple immunizations are simulated, the GC for the 2^nd^ and 3^rd^ rounds of affinity maturation are seeded with memory B cells output from the previous immunization. From experimental evidence it is known that the new GC is seeded by a variable number of B cells (tens to hundreds) [40]. In our simulations we used three, chosen at random from the pool of memory B cells. These are then allowed to proliferate in the dark zone without mutation until the B cell population reaches an initial size of 384 B cells (128 x 3). We note that the number of founding B cell clones is higher here than the number of B cells that were chosen to seed a GC upon the first immunization. Since we are not limited by the available experimental data, we increased this number to more accurately model the effects of clonal competition. Increasing this number much beyond three would be computationally intractable with the current model.

### Computing the breadth from the binding free energy

The breadth of a BCR is defined as the fraction of Ags in a panel that can be neutralized by the Ab. We compute the binding breadth *B*, which is based on the binding free energy ΔG of the BCR for all Ags in the panel:

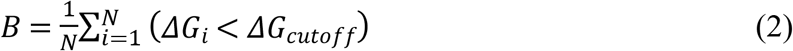

In this expression, *N* is the number of Ags in the panel, *ΔG_i_* is the binding free energy for the *i*-th Ag, and *ΔG_cutoff_* is an upper limit in the binding free energy above which the BCR is considered to be unable to bind to the Ag [46]. For the panel of Ags, we used the Seaman virus panel from the literature which is composed of 106 Ags [29]. To validate the binding free energy scoring function (see SI for additional validation details), we tested its ability to discriminate between the breadth of the mature VRC01 Ab from its putative germline VRC01GL Ab. This is of the utmost importance as the AM simulations are initiated from the germline BCR sequence, which then evolves over time into a more mature BCR sequence. We computed the binding free energy for VRC01 and VRC01GL for all 106 Ags in the Seaman panel and compared the two distributions (Fig. 5). For the calculations, PDB 5FYJ [28] was used as the template. From Fig. 5, we can see that the average binding free energy of the germline is shifted to higher values, indicating weaker binding, with a clear separation between the histograms for the mature and germline Abs. Using a cutoff binding free energy value of -9.9 kcal/mol, the computed breadth of the germline and mature Abs are 0.0 and 0.31, respectively. We note that lower thresholds lead to even greater separation between these values (e.g., a threshold of -9.7 kcal/mol leads to germline and mature Ab breadths of 0.0 and 0.64, respectively). However, -9.9 kcal/mol was found to be the best threshold to use to discriminate among the BCRs produced in the AM simulations, for which the binding affinity distributions were observed to vary widely depending on the temporal Ag administration protocol.

**Fig. 5.**
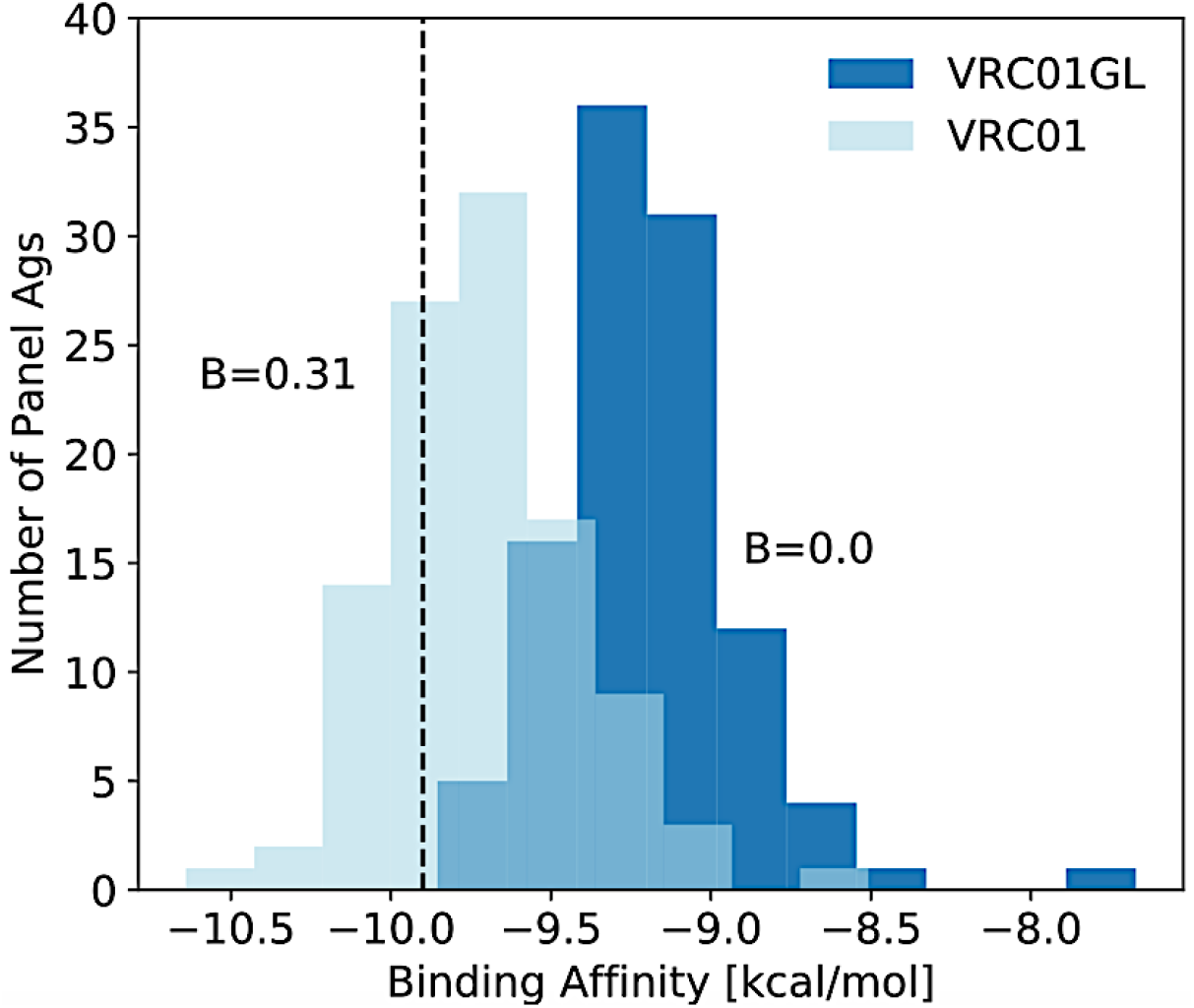
Binding free energy distributions of VRC01GL and VRC01 against a panel of 106 HIV antigens (Ags), using the scoring function developed in this work. A black dashed line indicates the threshold of -9.9 kcal/mol used to determine the breadth (B) of the BCRs. Note that the intermediate colored region arises from the overlap of the two distributions.

### *In silico* BCR (CD4bs) interfacial residue definition

In the crystal structure of VRC01 in complex with HIV gp120, nine residues were found to be in direct contact (within 4Å) with glycans on the surface of the Ag (Fig. 6A, right). Our scoring function for the BCR/Ag binding free energy was not parameterized to include the effect of glycans. Thus, we removed the glycan moieties from the structure files prior to the calculations and excluded these residues from our CD4bs interfacial Ab residue definition, which was used to determine the degree of interfacial composition matching of the BCRs from the AM simulations. We also excluded an additional two interfacial residues in VRC01GL that positionally aligned with these nine residues (Fig. 6A, left). All other interfacial residues identified for VRC01GL and VRC01 were considered for a total of 22 residues (see Table S2 in the SI). We note that inclusion of the 11 glycan-contacting residues in our calculations would lead to incorrect assessments of the degree of ICM achieved by the produced BCRs, because the driving force to evolve certain mutations against HIV glycans is not captured by our model. Despite the above residue exclusions, as shown in Fig. 6B, the set of 22 interfacial residues is sufficient to capture the major trends in composition matching observed in Fig. 3A, namely large increases in the basic and apolar interfacial fractions and a large decrease in the polar interfacial fraction. Amino acids were classified as follows: polar (S, T, C, Y, N, Q), apolar (G, A, V, L, I, M, W, F, P), acidic (D, E), and basic (K, R, H).

**Fig. 6.**
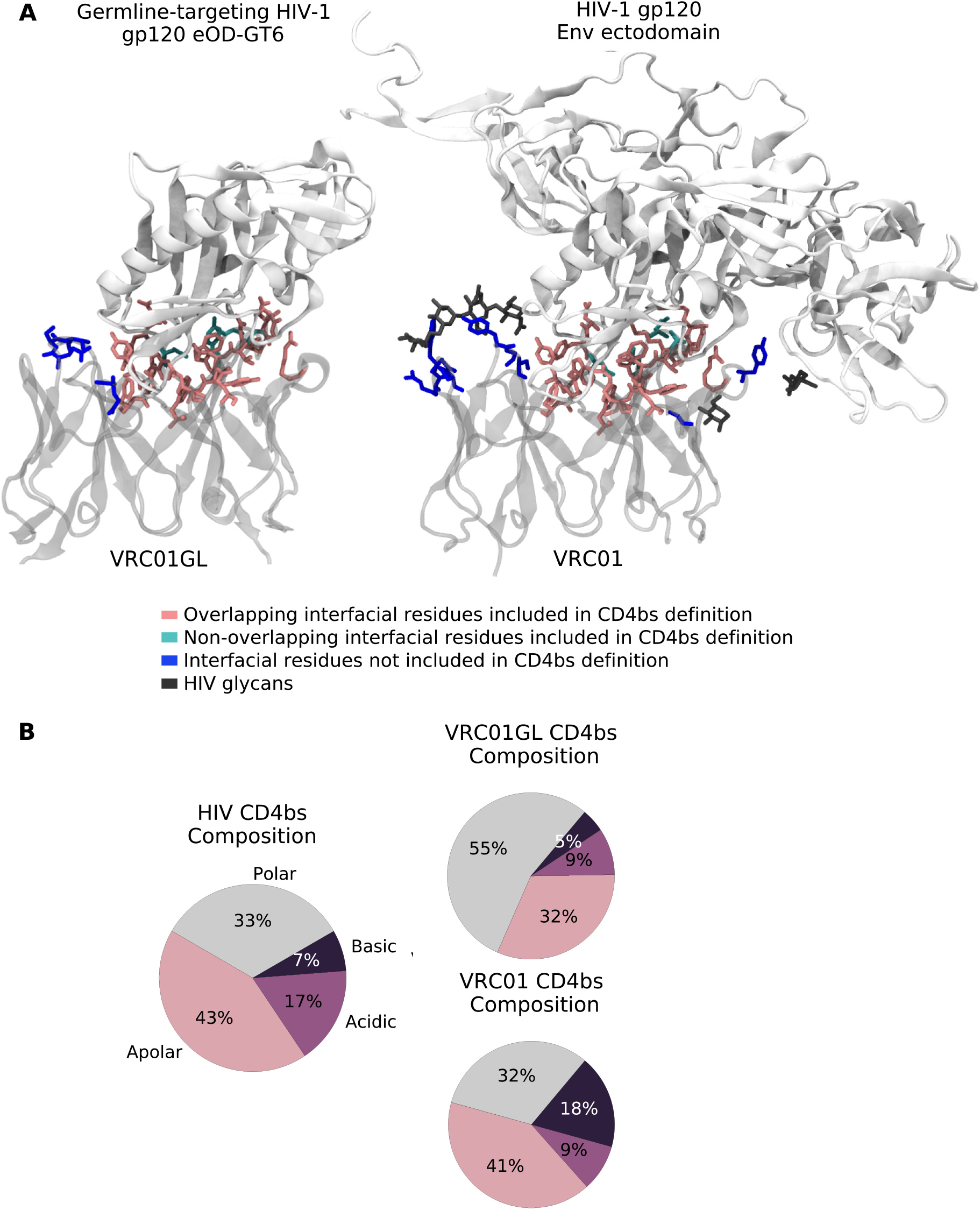
Definition of interfacial BCR residues used to characterize AM simulations. (A) Visual Molecular Dynamics (VMD [48]) snapshots of (left) VRC01GL and (right) VRC01, in complex with HIV-based Ags. Ags are colored in opaque silver and Abs in transparent silver. Residues included in the *in silico* CD4bs interfacial residue definition are colored pink if overlapping between the two Abs and colored cyan otherwise, and interfacial residues excluded from the definition due to contact or alignment with HIV glycans (gray) are colored blue. (B) Coarse-grained interfacial amino acid composition of the CD4bs of HIV (left), and of VRC01GL (top right) and VRC01 (bottom right) for the residues shown in pink/cyan in (A), which mimic the biological trends shown in Fig. 3A.

BCR interfacial residues in contact with conserved Ag residues were similarly identified using VMD. A total of 15 Ab residues were identified to be within 4Å of one of eight conserved Ag residues in PDB code 5FYJ (see Table S3 in the SI). These 15 residues represent 68% of the 22 total BCR interfacial residues listed in Table S2. The conserved Ag residues were identified by Conti et al. [20] as those for which the cost of mutational escape is high, according to the fitness landscape of HIV gp160 [47]. Consequently, these residues are generally highly conserved across many HIV strains and are commonly targeted by bnAbs against the CD4bs [20]. For each immunization of a given protocol, the number of BCR mutations that had arisen at one of the 15 identified residue positions was determined for each produced B cell clone to determine the extent of mutation towards conserved Ag sites. The results were then weighted by the size of each B cell clone and averaged across the two independent simulation trials.

### Interfacial composition/electrostatic pattern matching (ICM) calculations

For the ICM analysis of anti-CD4bs bnAbs (see Results), the following crystallographic structures were used: PDB codes 4JPK and 5FYJ for the germline [21] and mature [28] forms of VRC01, respectively, and PDB codes 5IGX and 3U7Y for the germline [30] and mature [31] forms of NIH45-46, respectively. Only the structure of the germline [30] Ab is currently available for 3BNC60 (PDB code 5FEC), whereas only the structure of the mature [32] Ab is available for CH103 (PDB code 4JAN). To overcome the challenge of missing structural data, we noted that the binding interface for VRC01 and NIH45-46 does not change significantly upon Ab maturation. Assuming that this holds for the other Abs, the same structure was used for the germline and mature forms of 3BNC60 and CH103 to identify interfacial residues in each case.

For the ICM analysis of the HIV CD4bs composition, interfacial Ag residues within 4Å of any Ab residue were identified from the crystal structures of multiple bnAbs in complex with HIV gp120, namely two structures of VRC01 (PDB codes 5FYJ [28] and 3NGB [49]), and one structure each of VRC18 (PDB code 4YDL [50]) and NIH45-46 (PDB code 3U7Y [31]). Sixteen common Ag interfacial residue indices were then identified across all of these data sets to isolate only the most conserved sites involved in Ab binding. These sites were used to determine the ICM of each of the 106 Ags on the Seaman virus panel [29], and the results were averaged across the panel.

The equation for calculating the ICM score for the BCRs in a single B cell clone *i* is shown below:

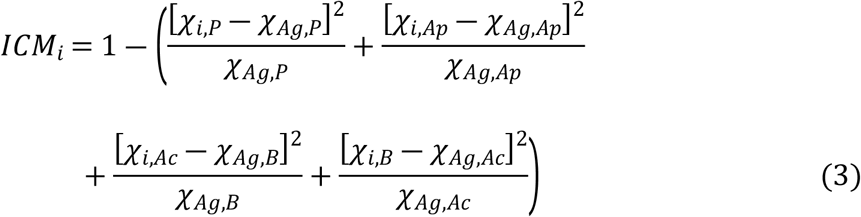

where *P*, *Ap*, *Ac*, and *B* stand for polar, apolar, acidic, and basic amino acid types, respectively, *χ*_*i*,*k*_ is the interfacial fraction of amino acid type *k* for clone *i,* and *χ*_*Ag*,*k*_ is the average interfacial fraction of amino acid type *k* for the Ag panel.

The ICM for an AM simulation is then defined as the average of the ICM scores of all individual clones (*ICM*_*i*_), weighted by the number of B cells in each clone (*N*_*i*_):

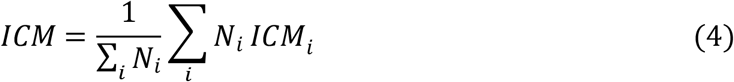

## Acknowledgements

Financial support was provided by Lawrence Livermore National Laboratory under Grant 17-ERD-043 (LLC Award B620960), by the Ragon Institute of MGH, MIT, and Harvard University (K.G.S and A.K.C), and by the CHARMM Development Project (M.K.).

## SUPPORTING INFORMATION

**Table S1.** Simulated vaccination protocols with 1, 2, or 3 sequentially administered Ags. Results are shown for the mean BCR breadth, mean degree of interfacial composition/electrostatic pattern matching (ICM), and the mean number of GC cycles. Standard deviations are on average ±0.05 for both mean BCR breadth and ICM, and ±2.6 for the number of GC cycles (see also Figs. 2 and 4 in the main text).

| # | Protocol | Mean Breadth | Mean ICM | Mean Cycles | # | Protocol | Mean Breadth | Mean ICM | Mean Cycles |
| --- | --- | --- | --- | --- | --- | --- | --- | --- | --- |
| 1 | WT | 0.00 | 0.75 | 7.5 | 19 | WT-HQ-EU | 0.92 | 0.84 | 34 |
| 2 | KR | 0.00 | 0.79 | 12 | 20 | WT-HQ-KR | 0.66 | 0.84 | 29 |
| 3 | HQ | 0.15 | 0.83 | 14 | 21 | KR-KR-KR | 0.97 | 0.79 | 68 |
| 4 | EU | 0.11 | 0.71 | 14 | 22 | KR-WT-HQ | 0.59 | 0.89 | 42 |
| 5 | WT-WT | 0.25 | 0.86 | 24 | 23 | KR-WT-EU | 0.67 | 0.83 | 54 |
| 6 | WT-KR | 0.63 | 0.86 | 26 | 24 | KR-HQ-WT | 0.26 | 0.75 | 46 |
| 7 | WT-HQ | 0.50 | 0.83 | 24 | 25 | KR-HQ-EU | 0.88 | 0.84 | 55 |
| 8 | KR-KR | 0.55 | 0.92 | 38 | 26 | HQ-KR-WT | 0.38 | 0.88 | 30 |
| 9 | KR-WT | 0.39 | 0.84 | 32 | 27 | HQ-KR-EU | 0.76 | 0.89 | 49 |
| 10 | KR-HQ | 0.57 | 0.73 | 42 | 28 | HQ-EU-WT | 0.82 | 0.85 | 36 |
| 11 | HQ-KR | 0.25 | 0.88 | 21 | 29 | HQ-EU-KR | 0.83 | 0.88 | 36 |
| 12 | HQ-EU | 0.25 | 0.88 | 21 | 30 | EU-EU-EU | 0.92 | 0.83 | 51 |
| 13 | EU-EU | 0.60 | 0.82 | 25 | 31 | EU-WT-KR | 0.95 | 0.84 | 38 |
| 14 | EU-WT | 0.12 | 0.71 | 18 | 32 | EU-WT-HQ | 0.78 | 0.8 | 42 |
| 15 | EU-KR | 0.13 | 0.72 | 22 | 33 | EU-KR-WT | 0.67 | 0.81 | 40 |
| 16 | EU-HQ | 0.49 | 0.76 | 24 | 34 | EU-KR-HQ | 0.88 | 0.88 | 39 |
| 17 | WT-WT-WT | 0.53 | 0.86 | 30 | 35 | EU-HQ-WT | 0.42 | 0.77 | 30 |
| 18 | WT-KR-EU | 0.85 | 0.86 | 32 | 36 | EU-HQ-KR | 0.82 | 0.81 | 36 |

**Table S2.** List of 22 amino acid residues used in the *in silico* CD4bs interfacial BCR residue definition and Fig. 7 (main text). Residue numbers and identities correspond to those in PDB codes 5FYJ^1^ for VRC01 and 4JPK^2^ for VRC01GL.

| # | Residue | VRC01GL | VRC01 | # | Residue | VRC01GL | VRC01 |
| --- | --- | --- | --- | --- | --- | --- | --- |
| 1 | 33 | Y | T | 12 | 60 | A | A |
| **2** | 47 | W | W | **13** | 61 | Q | R |
| **3** | 50 | W | W | **14** | 64 | Q | Q |
| **4** | 52 | N | K | **15** | 71 | R | R |
| **5** | 53 | N | R | **16** | 99 | D | D |
| **6** | 54 | S | G | **17** | 100 | Y | Y |
| **7** | 55 | G | G | **18** | 100 | N | N |
| **8** | 56 | G | A | **19** | 100 | W | W |
| **9** | 57 | T | V | **20** | 91 | Y | Y |
| **10** | 58 | N | N | **21** | 96 | E | E |
| **11** | 59 | Y | Y | **22** | 97 | F | F |

**Table S3.**
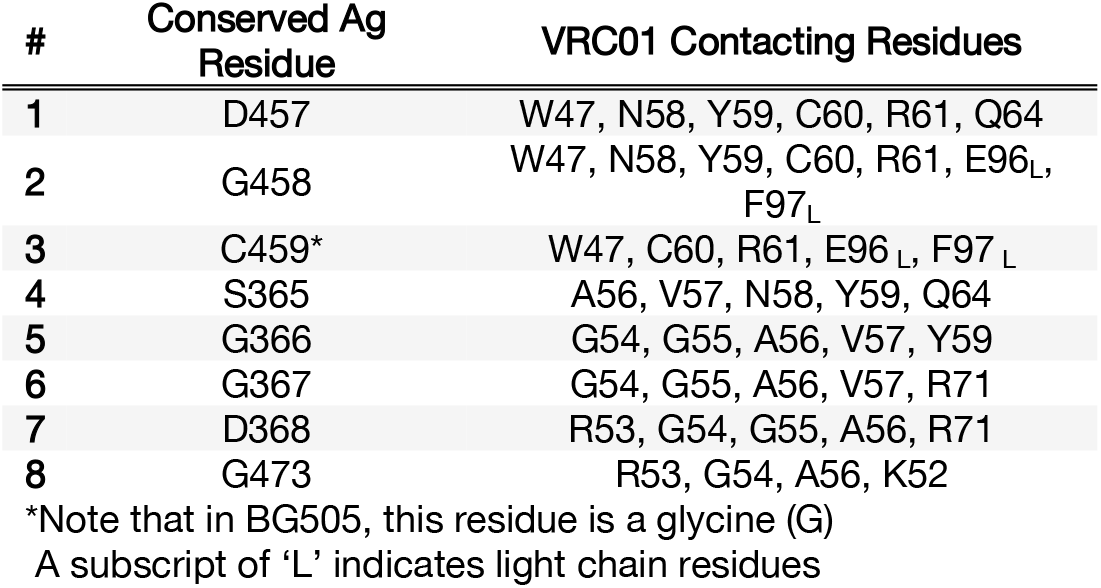
List of conserved residues in the CD4bs of HIV, as determined by Conti et al.^3^, and the corresponding residues of VRC01 in contact with the Ag residues, used to characterize BCR conserved site binding (Fig. 2D, main text). Residue numbers and identities correspond to those in PDB code 5FYJ^1^.

**Fig. S1.**
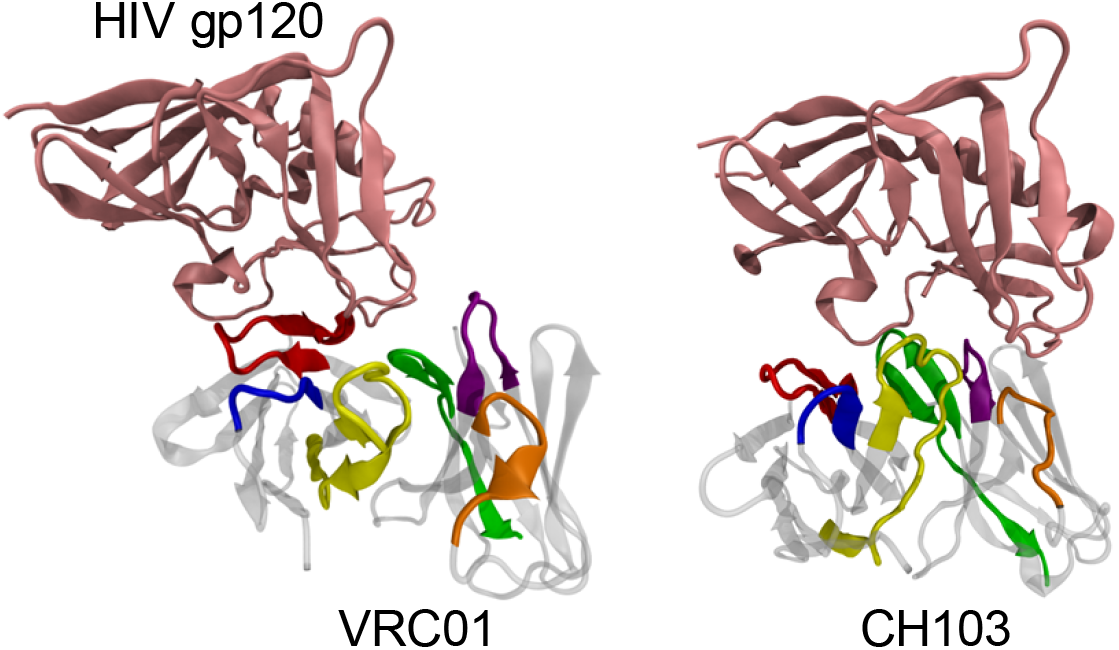
Snapshots from Visual Molecular Dynamics (VMD^4^) of VRC01 (left) and CH103 (right) in complex with gp120-based Ags. Ags are shown in pink and Abs in transparent gray, with the six complementarity-determining regions (CDRs) colored in red (CDRH2), orange (CDRL2), yellow (CDRH3), green (CDRL3), blue (CDRH1), and purple (CDRL1).

### Four key biological principles dictate the types of observed mutations

Our results establish interfacial composition/electrostatic pattern matching as a major biological driving force in the affinity maturation trends of anti-CD4bs bnAbs. To gain insight into other factors that might influence the evolutionary trajectories of these Abs, we examined in more detail the specific interfacial mutations that VRC01GL made upon maturing into its bnAb form. We considered 33 interfacial residue indices in this analysis, including the 21 overlapping residue indices between VRC01GL and VRC01, as well as the 4 additional residue indices identified solely in VRC01GL and 8 additional residue indices identified solely in VRC01. Of these 33 residues, 14 were mutated during the evolution of VRC01GL into VRC01. We characterized each of these 14 mutations according to which amino acid (AA) type it was mutated from/to in VRC01GL and VRC01, respectively, and then determined the overall fraction of mutations from/to each AA type. These results are presented in Fig. S2A as a heat map.

In line with the results of Fig. 3A (main text), the results of Fig. S2A show that a significant fraction of the interfacial mutations was made away from polar AAs. This is true even considering the large fraction of mutations made away from polar AAs simply to other polar AAs. We sum up the fraction of mutations made away from polar AAs (Fig. S2A, top row) and subtract this from the summed fraction of mutations made to polar AAs (Fig. S2A, far left column), to see that the overall selection tendency for polar AAs at the interface, or μ_*Polar*_, is strongly negative (-0.50). Similarly, we find a strong positive selection tendency towards apolar (μ_*Apolar*_ = +0.15) and basic (μ_*Basic*_ = +0.28) AAs at the interface, also in line with the results of Fig. 3A (main text). However, the same analysis shows a mild positive selection tendency towards acidic residues (μ_*Acidic*_ = +0.07), seemingly at odds with the results of Fig. 3A. In VRC01GL, 3/25 interfacial residues are acidic, compared to 3/29 residues in VRC01, resulting in the decreased interfacial acidic fraction in Fig. 3A. For one of the acidic interfacial residues in VRC01, its identity in VRC01GL was an apolar residue, explaining the observed tendency to mutate towards acidic residues in Fig. S2A.

These results raise a series of questions, such as why the need for increasing the basic interfacial fraction was met primarily through making mutations away from polar residues versus away from apolar residues, or why so many seemingly unessential polar-to-polar and apolar-to-apolar transitions occurred. Answers to these questions are in part obtained by considering the average number of nucleotides required to transition from one type of AA to another. These values are shown in Fig. S2B and were obtained by calculating the average pairwise distance between all codons encoding for each pair of AA types. From Fig. S2B, we can see that fewer nucleotides are required on average to make a polar-to-basic transition than to make an apolar-to-basic transition. Also, given a certain AA, the data shows that it is easiest to mutate to a different AA in the same category, due to the similarities in their codons. This explains, at least in part, the many polar-to-polar and apolar-to-apolar mutations that were made during the evolution of VRC01. Another contributing factor for these mutations is that many polar-to-polar and apolar-to-apolar mutations are “neutral” mutations, which are neither positively nor negatively selected during AM. It is known that VRC01-class Abs typically emerge only after many years of infection^5,6^, and so the Abs have long maturation periods over which to accumulate these largely irrelevant mutations. The reasons for the lack of basic-to-basic and acidic-to-acidic mutations – or any mutations away from charged AAs, for that matter – are two-fold. We can observe from Fig. 3A (main text) that charged AA types make up the lowest interfacial fractions by far in VRC01GL and thus they are much less likely to be targeted for mutation, simply from a probability standpoint.

In addition, nucleotides in codons encoding for polar AAs are also, on average, more mutable than those in codons that encode for the other AA types (Fig. S2C). The values reported in Fig. S2C were determined using the model developed by Yaari *et al*.^7^ (see Methods in main text). We arrived at the values reported in Fig. S2C by computing the average mutability score of all 5-mers containing a given codon for all codons encoding for a particular AA type. Computing the mutability scores for the individual nucleotides in the CDRs of VRC01GL in a similar manner, Fig. S2D shows that there are many more nucleotides in codons that encode for polar AAs – and even apolar AAs – with high mutability scores, than there are basic or acidic AAs with high mutability scores (e.g., 17 data points classified as polar have a score > 2, versus 6, 2, and 1 data points classified as apolar, basic, and acidic, respectively).

To summarize, four key biological principles dictate the types of observed mutations in VRC01 as compared with VRC01GL. These are the mutations that we expect to observe in our simulations. From a targeting standpoint, both the initial, relative proportions of different AA types at the interface (principle #1) and the mutability of the codons encoding for the interfacial AAs (principle #2), have a large impact on the evolved mutations.

Moreover, mutational patterns are heavily influenced by the difficulty of transitioning from one AA type to another, which determine the length of time required for the GC reactions to take place (principle #3). Lastly, from a selection standpoint, a large initial difference between the interfacial AA composition of the germline BCR and that of the CD4bs, creates a strong driving force to evolve mutations that lead to increased interfacial composition/electrostatic pattern matching (principle #4). The degree of correlation between these principles and the effects that changes in the parameters have on the ensuing number and type of acquired Ab mutations will be explored in detail in future studies (e.g., the impact on AM of using germline sequences with different initial mutability patterns or degrees of interfacial composition matching). Here, we explore the impact of administering multiple variant Ags in different temporal patterns on the ability of the BCR population to evolve mutations that (1) lead to high breadth, and (2) lead to an increased degree of interfacial composition/electrostatic pattern matching against the CD4bs.

**Fig. S2.**
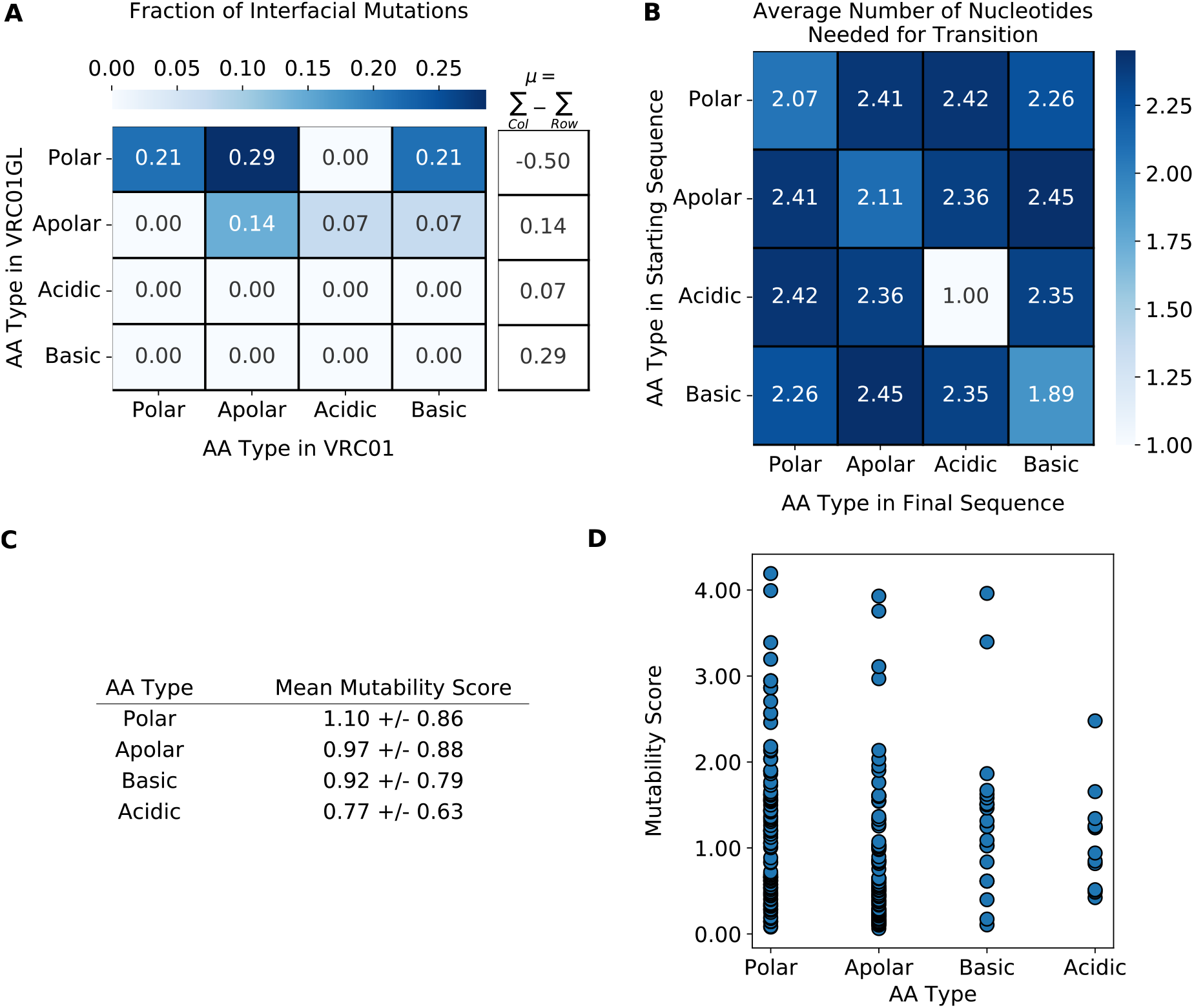
Basic biological principles govern the mutations acquired by VRC01 during AM. (A) Heat map of the fraction of interfacial mutations (n=14) made from each amino acid (AA) type in VRC01GL to each AA type in VRC01. The overall mutation tendency for each AA type, μ, is computed as the sum of the relevant column values subtracted by the sum of the relevant row values. (B) Heat map of the average number of nucleotides needed to transition from one AA type to another. (C) Mean mutability of the different AA types, based on the model of Yaari *et al.*^7^. (D) Mutability score of each nucleotide in VRC01GL, grouped and colored by the type of AA encoded by the codon to which the nucleotide belongs.

**Fig. S3.**
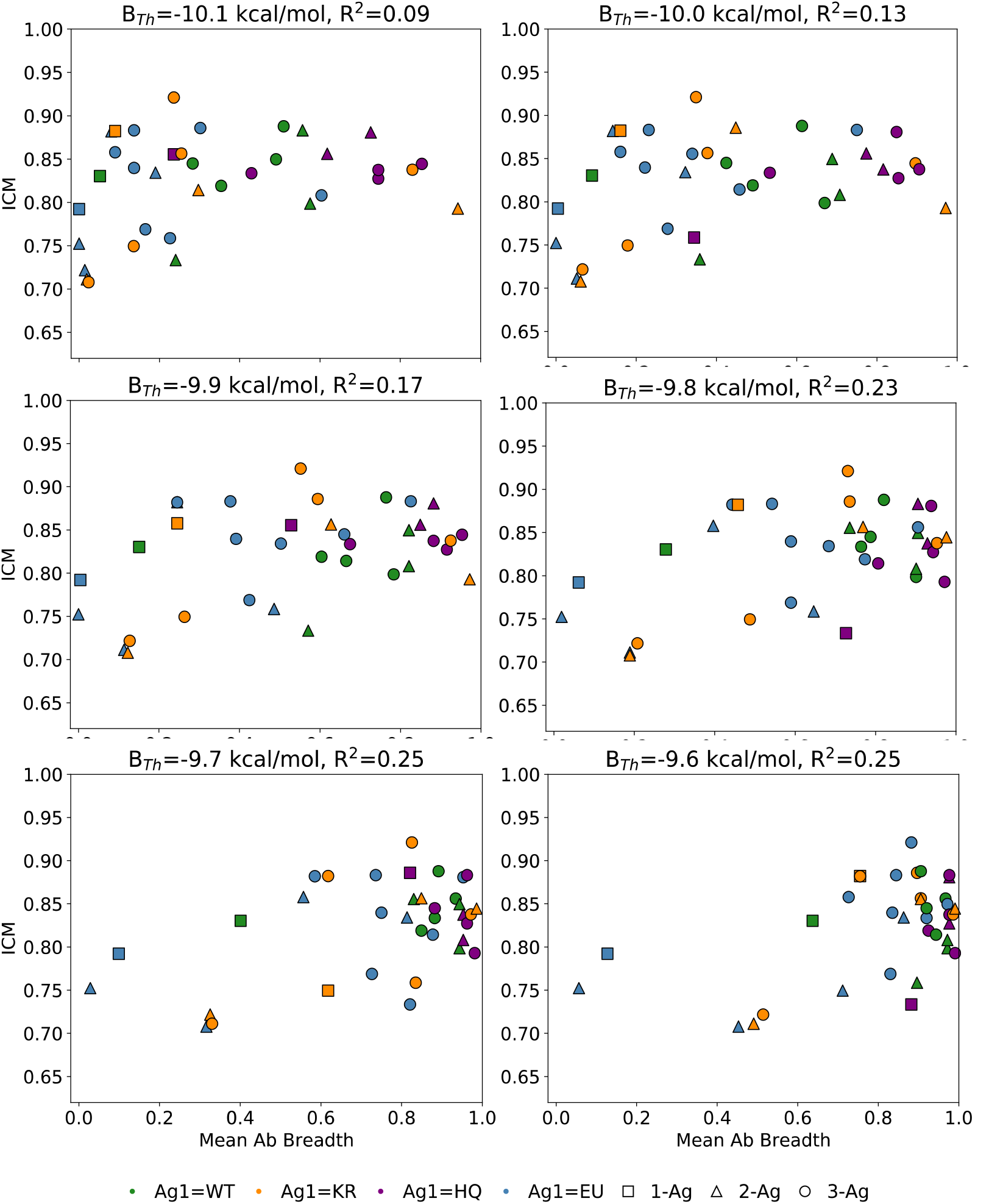
Mean BCR breadth of vaccination protocols with t=1, 2, and 3 single-Ag sequential immunizations versus the mean weighted degree of interfacial composition and electrostatic pattern matching (ICM), computed using different thresholds for determining the BCR breadth. Error bars are only shown for the degree of ICM for clarity and represent the standard deviation of two independent simulations of n=1,000 GC reactions.

### Analysis of mutational trajectories of individual clonal BCR sequences

The results for the EU-based protocols (Fig. S4A) and HQ-based protocols (Fig. S4D) were discussed in the main text. For the KR-based protocols (Fig. S4B), we find that many B cell clones evolved basic-to-polar mutations after just the first immunization with KR, and this trend continued after a second immunization with KR (Fig. S4B, blue line), shedding light on the high ICM score of the KR-KR protocol, as mentioned earlier. However, after a third immunization with KR, we observe a reversion back to the polar and basic interfacial fractions – and the lower ICM score – obtained after the first immunization with KR. For the rest of the KR-based protocols, no clear trends exist in the types of mutations that are evolved after the second and third immunizations, leading to the wide array of final ICM scores reported in Fig. 4 (main text).

For the WT-based protocols (Fig. S4C), the first immunization resulted in little change in the interfacial amino acid fractions compared to VRC01GL. After the second immunization, regardless of which Ag was administered, we observe relatively large decreases and increases in the polar and basic interfacial fractions, respectively. However, these ICM gains are offset by mutations that decreased the apolar interfacial fraction for all four WT-based protocols. The third and final immunization, again regardless of which Ag was administered, generally led to little change in the interfacial fraction of any amino acid type, resulting in the intermediate ICM scores for the 2- and 3-Ag WT-based protocols shown in Fig. 4 (main text).

**Fig. S4.**
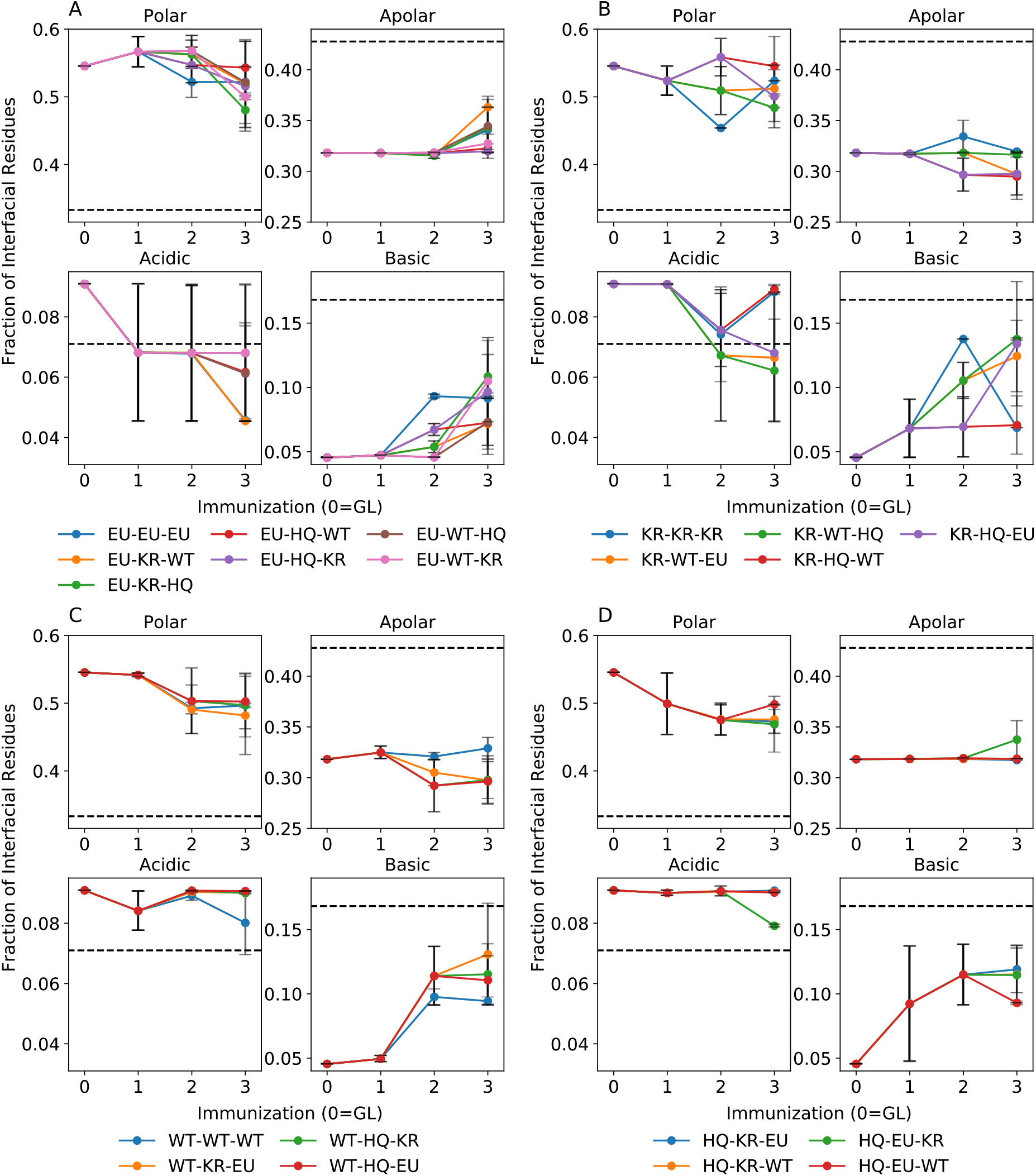
Fraction of different aa types at the BCR/Ag interface in BCR sequences produced after administration of protocols beginning with (A) the EU Ag, (B) the KR Ag, (C) the WT Ag, or (D) the HQ Ag. Black dashed lines indicate the interfacial amino acid fractions in the CD4bs of HIV. “Immunization 0” refers to the GL-targeting scheme that we assume takes place prior to vaccination (see main text).

### Modeling the B cell receptor/antigen binding free energy

The objective is to find a protocol to compute binding affinities between a given arbitrary sequence of an antibody and an HIV antigen that would require the minimum computer time. At each cycle of the affinity maturation simulation this protocol will be called hundreds of times, so it is imperative to reduce the computational time as much as possible. The methods we use are based on all-atoms descriptions of the antibody-antigen protein-protein complex, so the first task is to generate a reasonable model of the complex. We found that Modeller^8^ is the quickest and most reliable method to produce such models, given that an initial template of a related system is available. As templates we use crystal structures of HIV antibodies bound to the gp120 HIV surface protein. The first step of our protocol is thus to create one model of the full complex with Modeller, by grafting the target sequence onto the template. We chose to use a very quick refinement protocol in order to reduce the computation time. Hydrogen atoms are not included in this step, since it was found that in this quick protocol, they often generate the wrong chirality, so only the positions of heavy atoms are generated. The second step is to optimize the structure. We need to add hydrogen atoms, check for the presence of disulfide bonds, and check for the presence of errors in chirality, in particular around the alpha carbon of each protein residue. Last, the system is energy minimized with 10 steps of steepest descent (SD), followed by 10 steps of adapted-based Newton-Raphson (ABNR). This is expected to fix all mayor problems in the structure, in particular to remove atomic clashes. These checks were all performed using CHARMM^9^. Then the model of the complex is split, to create a model of the ligand (the antibody) and the receptor (the antigen). To compute the binding affinity we use the Rykunov-Fiser statistical pair potentials (RFSPP)^10^. These potentials compute the stability of each protein (G_x_), and the binding affinity (dG) can be computed as the difference dG = G_cpx_ – G_lig_ – G_rec_. This whole procedure was repeated 12 times starting with the creation of the model with Modeller using different initial random seeds. The final RFSPP scoring was obtained as the average over the 12 values. Two RFSPP are available: an all-heavy atoms model (RF_HA_SRS) and a beta-carbon directional potential (RF_CB_SRS_OD). While the second is slightly faster, the first produces higher quality results. These scores are not directly usable as binding affinity values and need to be rescaled to better fit experimental data. The described protocol requires about 12 minutes of computer time to compute one binding affinity (about 1 minute per model), but the models can be run in parallel, requiring only 1 minute of user time on a 12-core machine.

To validate the accuracy of the RFSPP scoring, the computed values were compared with 106 experimental binding affinities of HIV antibody/antigens complexes from the literature^11^. Data were collected for four complexes: the resurfaced stabilized core 3 (RSC3) antigen bound to antibodies VRC01, VRC03, and VRC-PG04, and antigen 93TH057 bound to antibody VRC01. For these four complexes experimental data are available for simple single point mutations and, in the last case, also for few insertion and deletions. For each available experimental dG, the RFSPP score was evaluated as described, and the obtained scores were compared with the experimental dG. As templates we used PDBs 3NGB^12^, 3SE8^13^, 3SE9^13^, and 5FYJ^1^, respectively, for the four base complexes. The scale and unit of measure of the RFSPP score is uncertain, so a linear regression is performed to find the best parameters (m and q) to interpret the scores as binding affinities in kcal/mol: dGscore = m * RFSPP + q. The whole set of experimental binding affinities was split in half randomly: the first half was used to fit m and q by the linear regression and the second half was used to evaluate the fit, specifically the Pearson (Rp) and Spearman (Rs) correlation coefficients, the Mean Absolute Error (MAE) and Root Mean Square Error (RMSE). Due to the use of a random split in the training and validation sets, the training can be repeated a number of times and different models can be obtained. To reduce this randomness and avoid picking an arbitrary initial random seed, the full training was repeated 10000 times and all Rp, Rs, MAE and RMSE were stored. To remove all solutions with obviously bad properties (like very low Rp, or very high MAE) a Pareto optimization strategy^14^ was adopted. It selects only the models for which no other model exist that have both higher Rp and lower MAE. This procedure constructs a Pareto front that contains only 4 models. Sorting the models in the Pareto front, the first and the last are the ones that optimize the Rp and MAE, respectively. Seeking a model that is balanced between the two limits, the third model of the front was chosen. The obtained linear fit with the RF_HA_SRS statistical pair potential shows a Rp=0.74 and Rs=0.74, while the MAE and RMSE are 0.50 kcal/mol and 0.59 kcal/mol, respectively. Fig. S5 shows the obtained regression. The main negative feature is the low slope of the linear regression, which indicates that the computed dG will be compressed in a smaller range of values with respect to the expected experimental values. This problem is common to all tested linear regressions, even using very different descriptors (other statistical potentials, implicit solvents, buried surface area based, etc.). The final linear regression result is dG_score_ = m * RFSPP + q, with m=0.00619 and q=-8.174. A survey of similar parameters showed qualitatively very similar results.

**Fig. S5.**
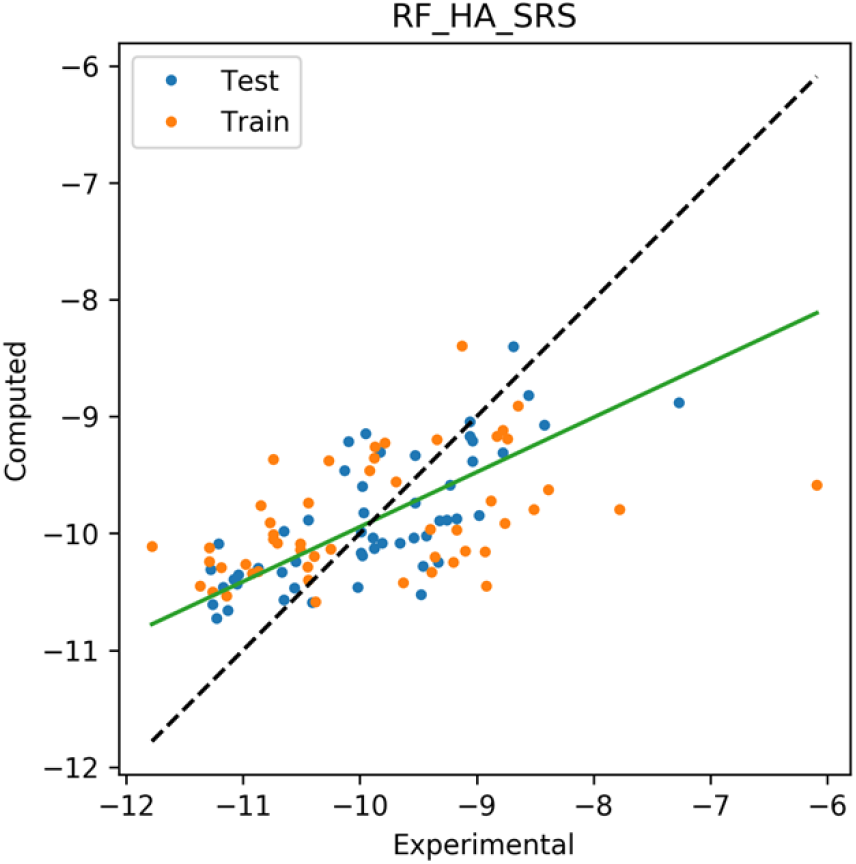
Linear correlation between experimental and computed binding affinities using the RF_HA_SRS statistical pair potential. Each dot represents one binding affinity value. The data in orange are used for the training of the linear regression, while in blue for the validation.

### Scoring function validation: scan for spurious mutations

The affinity maturation protocol involves the stepwise accumulation of mutations from the germline to move towards some mature-like antibody. Intuitively, we expect that a single point mutation cannot produce a too large gain in binding affinity, and also that there are no particularly good mutations that always increase the binding affinity. Due to the use of a very crude and fast scoring function, we need to explicitly check for these cases in order to avoid strong artifacts in the simulations. For these reasons we identified 25 residues around the binding site of the VRC01 antibody bound to the BG505 SOSIP antigen and we mutated each of those residues from the germline sequence into each possible amino acid (19 possibilities, excluding the original one), and computed their binding affinity to three antigens: BG505 and the antigens CH115_12 and MW965_26 from the Seaman panel. Fig. S6a shows the range of all 475 (= 25*19) relative change in binding affinity, sorted by magnitude for the three antigens. The highest favorable and unfavorable changes are of only ±2kcal/mol, with the vast majority of mutations being neutral (close to zero). This is the expected behavior for single point random mutations. With the same data, it is also possible to check if some mutations (e.g., mutation to cysteine) are always beneficial mutations. This would imply that a bias towards that particular amino acid is present in the scoring function. The bar plot in Fig. S6b shows the probability that mutating to each amino acid results in an improved binding affinity (i.e., an amino acid with a probability of one means that while scanning the 25 residues in the binding site, mutating any of those resides to that amino acid results in a better binding). Intuitively, we expect that no “preferred” amino acid should exist, which means that the probability should never be close to one, but “terrible” amino acids, with probability close to zero, could exist, e.g., charged amino acids, which are much more sensitive to their environment. The results obtained from antigen BG505 are in good agreement with what is expected, with no preferred mutations (max probability is about 0.6 for W, Y, and H which are all aromatic) and there are few “terrible” amino acids (D, E, K, and Q, all charged or polar). Antigens CH115_12 and MW965_26 behave similarly.

**Fig. s6.**
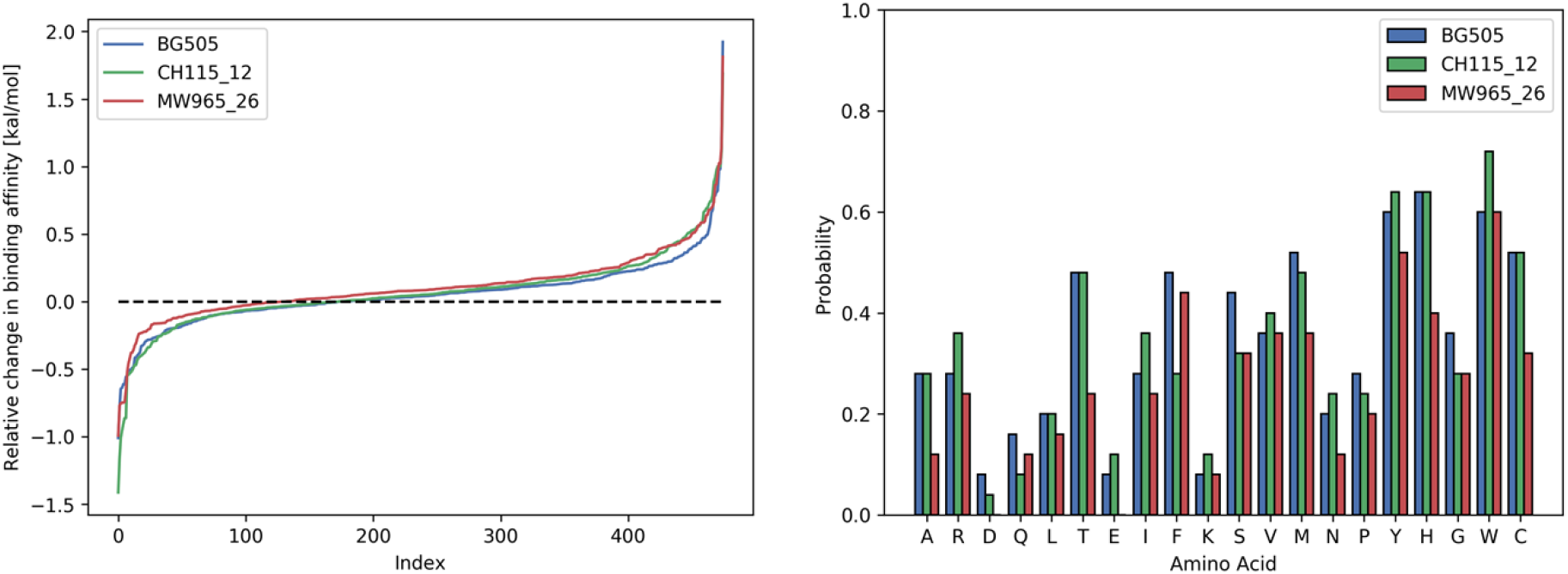
a) Relative binding affinity changes while mutating the 25 residues of VRC01GL at the binding site into each possible amino acid, sorted by magnitude. The range of the magnitudes is as expected limited with most mutations neutral (close to zero). b) Bar plot of the probability that mutating to a given amino acid improves the binding affinity. There are no preferred amino acids with very high percentages.

